# INFLUENCE OF SOME LEGUMINOUS TREES AND ARBUSCULAR MYCORRHIZAL FUNGI ON THE PHYSIOLOGY AND YIELD OF DIOSCOREA ROTUNDATA -POIR

**DOI:** 10.1101/2020.08.16.252908

**Authors:** Ezekiel Taiwo Afolayan

## Abstract

This work compares the physiological and yield characteristics of white yam (Dioscorea rotundata – Poir) under Arbuscular mycorrhizal fungi (AMF) inoculation, green manures of Gliricidia sepium, Leucaena leucocephala and other soil amendments. The experiment was conducted on the plot of land that had been overcropped, located at the back of the male Hostel, Federal College of Education, Abeokuta, Ogun State, Nigeria. The land was cleared and heaped at 1m x 1m apart. The experimental design employed was a complete randomized design in 5 replicates. The treatments were Glomus deserticola (GD), Glomus fasciculatum (GF), Gliricidia sepium (GS), Leucaena leucocephala (LL), Poultry manure (PM) and NPK fertilizers. Soils were dug from the heaps, 20 g of the inoculums of AMF (GD/ GF) were poured into the dug hole, seeds were laid on it and covered with soil (for GD & GF treatments). Others were applied at one week after sprouting. Growth and yield Parameters were determined at harvest while relative water and chlorophyll contents were measured forth nightly from 10 weeks after treatment. Data obtained were subjected to ANOVA while means were separated by Duncan multiple range test at P> 0.05. Results showed that growth, yield and physiological characters were enhanced in GD, GD+GF, GS and PM treated plants more than in inorganic fertilizers treated plants. There was a positive significant relationship between white yam’s growth, physiology and tuber yield. The study justifies the use of plant/animal manures and Arbuscular mycorrhizal fungi in place of inorganic fertilizers.

## INTRODUCTION

White yam (*Dioscorea rotundata* – Poir) account for more than 200 dietary calories per person each day for an estimated 60 million people, especially in the yam producing countries like Nigeria, cote d’lvoire, Ghana and Cameroun (Zannou 2004; Zannou *et al*., (2007). Nigeria is the largest producer of yam (IITA, 1993; FAOSTAT, 1997; Aighewi *et al*., 2002). Economically yam provides job for both the farmers and the women who are marketers of the products. It is also rich in minerals sources of proteins, fats and vitamins, yam is used as a foaming agent in brewing industries, manufacturing of drugs, insecticides, flavours, fragrances, drinks and cosmetics. It also plays a significant role in the cultural life of rural communities in Benin, Igbo and Yoruba people of Nigeria (Onwueme, 1978; Yadana, 2009).

Arbuscular mycorrhizal (AM) fungi are mutually beneficial microorganisms that form a symbiotic association with the roots of vascular plants. The host Plants supply the Arbuscular Mycorrhizal fungi with sugars, while the fungi scavenge for immobile nutrients, e.g. Phosphorus. Mycorrhizal associations are functionally diverse both in effectiveness of the fungi as symbionts and in responsiveness of the plants in terms of total P uptake, growth, and/or reproduction (Read, 1998; Van der Heijden, and Sander, 2002; Jakobsen, 2002). Nitrogen and phosphorus deficiency are common in soils in the tropical and subtropical regions. This constitutes the main limiting nutrients in semi-arid and dry sub-humid Savanna (Bationo *et al*., 1992: Buerkert *et al*., 2001). Sequel to this development, farmers in these regions, especially in Africa, employed the use of inorganic fertilizers which may not be ecologically friendly.

Arbuscular mycorrhizal fungi are important and will be of tremendous assistance in sustainable agriculture in Africa because they improve plant water relations and thus increase the drought resistance of host plants (Allen and Allen, 1986; Naidoo, 1986), and they increase mineral uptake, which reduces the use of fertilizers. Improved plant water status and changes in water relations have been attributed to a wide variety of mechanisms (Smith, and Gianinazzi-Pearson. 1988; Davies *et al*., 1992).

Woody legumes are important for re-vegetation of ecosystems that have low amounts of available N and P in order to mitigate the effects of deforestation and maintain sustainable land-use management Danso *et al*., 1992). Leguminous trees are nitrogen fixing trees (NFTs) which helps in the restoration and/or maintenance of soil fertility. Some of the examples of these NTF are *Gliricidia sepium and Leucaena leucocephala* which are fast-growing perennial leguminous trees for soil nutrient enrichment, N2-fixing capability and P assimilation and arbuscular mycorrhizal (AM) fungi which lives in the roots of plants as symbiotic organisms that helps the plants to absorb nutrients especially Phosphorus (Habte and Turk, 1991; Liyanage *et al*., 1994).

Macro and micronutrients together with some soil micro and macro organisms are both required for normal growth and development in plants. These, in their right proportion, are the major factors for enhanced yield and production of food. There is increase in plant metabolites as a result of the application of organic manure, this enhanced metabolic activities leading to improved growth characters. This increase in growth characteristics (i.e. stem, leaves, and roots) leads to higher photosynthetic activities and accumulation of higher photo assimilates (Yadana *et al*., 2009).

The use of organic fertilizers helps in preserving the ecosystem, maintain soil fertility by supplying of balanced nutrients, decomposition of harmful elements, improves soil structure, root development, and makes soil water available to plants roots and promote plants productivity. Organic manure increases soil mineral elements, crop productivity, plants’ health and mineral utilization efficiency (Kaur *et al*., 2005; Murmu *et al*., 2013). Green manure of leguminous plants enhanced the stem diameter, leaf number, leaf area, pod length, weight and diameter of edible fresh Okra plants (Benjawan *et al*., 2007).

The objective of this work was to examine the growth and yield promoting abilities of arbuscular mycorrhizal fungi, green manure of *Gliricidia sepium, Leucaena leucocephala* and other soil amendments in an overcropped soil.

## MATERIALS AND METHODS

This trial was carried out in 2017 and 2018 at the nutrients-depleted plot of land behind the male students’ hall of residence at the Federal College of Education, Abeokuta, Ogun State, South-west Nigeria. The land was cleared, loosened and heaped at 1m^2^ x 1m^2^ apart. Twenty grams (20 g) of mycorrhizal fungi inoculums (*Glomus deserticola* and *Glomus fasciculatum*) were measured into the dug hole on the heap and the seed yams (450 – 500 g) were placed on the inoculums before covering with the soil in AMF treatment plots. Seed yams were planted in the heap and mulched with 800 g *Gliricidia sepium* and *Leucaena leucocephala* as applicable. Poultry manure (80 g) and NPK fertilizer (40 g) was applied at one week after sprouting for poultry manure and NPK treatments. Staking was done with one stake per stand. Weeding and other agronomic practices were done as when due.

Plants performance was assessed through the following parameters:

a. **Shoot and Root weight**: These were determined at harvest. Plants parts (shoot and roots) per plant were carefully collected and weighed.
b. **Tuber diameter and length**: The diameter and length of the tubers per plant were determined using meter rule immediately after harvesting.
c. **Tuber weigh:** At harvest, tubers of yams per plant were placed on the weighing balance for tuber’s weight on the field.
d. **Chlorophyll content of leaves:** This was done to determine the health of the plants. It was determined using pocket chlorophyll meter MO-SPC-SPAD 502 (Konica Minolta, USA) because it was done on the field. Leaves of each treatment were inserted into the machine while on the field, the chlorophyll content per plant is displayed on the screen of the chlorophyll meter.
e. **Relative water contents**: Fresh leaves were collected from each treatment, each leaf was cut with cork borer into small discs, which were weighed and recorded (sample’s Fresh Weight - W). The samples were then hydrated to full turgidity in distilled water for 4 hours under normal room conditions. These were removed from the water after 4 hours and the adhering moisture was quickly dried off with filter paper and immediately weighed (Turgid Weight - TW). Samples were then dried in an oven at 80°C for 24 hours, cooled in desiccators and later weighed (the dry weight-DW).

The following formula was used to determine the Relative Water Content (R W C).

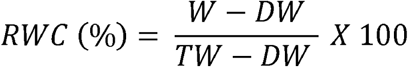

Where W = sample fresh weight; DW = sample dry weight; TW = sample turgid weight (Dingma, 2002).

Data obtained were subjected to statistical analysis using ANOVA (SPSS version 20), while treatments means were separated with Duncan multiple range test at 5% level of probability.

## RESULTS AND DISCUSSION

Chlorophyll and relative water contents of white yam under AMF, green manure of GS, LL and other soil amendments were determined and results were represented on figures 1and 2. Chlorophyll synthesis was enhanced by the application of GS, LL, GE, GM, PM and combined treatments of all the organic manures and arbuscular mycorrhizal fungi. A higher amount of Relative water content was observed in white yam treated with LL, PM, GF, GD, GS together with their combined treatments. There was a significant relationship between the water content and the chlorophyll contents synthesized in organic manure treated (GS, LL, GF, GD, PM in all organic manure treated plants (GS, LL, GD, GF, PM, GD+GF, GM+LL, GM+PM, GM+GS, LL+GF, LL+PM and LL+GS) when compared to the inorganic fertilizers and the untreated white yam. This may be due to increased absorption of water, N, and P in AMF (GM and GF) inoculated plants and readily absorbable nutrients made available by the organic fertilizers which are responsible for normal metabolic activities in plants (Changxian *et al*., 2008). This result corroborates the report of some workers who reported an enhanced chlorophyll and relative water contents in white yams treated leguminous combined with AMF (Afolayan, 2017; Oyetunji, & Afolayan, (2019).

Growth and yield characters of white yam treated with Arbuscular mycorrhizal fungi (AMF), green manure of *Gliricidia sepium, Leucaena leucocephala* and other soil amendment were correlated with each other and the results are presented in Table 3. It was observed that there was positive relationship between the shoot weight and the root weight (r= 0.655); tuber weight (r = 0.677); tuber diameter (r= 0.563) and tuber length (0.732) at P>0.05 level of significance. Root weight had a significant and positive correlation with tuber weight (r=0.850) at p> 0.05 level of significance.

**Table 1:** Chlorophyll contents (μmol/m^2^) of Dioscorea rotundata treated with AMF, green manure of *Gliricidia sepium, Leucaena leucocephala* and other soil amendment under continuous cropping systems

| Treatment | Chlorophyll contents($\mu\text{mol}/\text{m}^2$) | | | | |
| --- | --- | --- | --- | --- | --- |
|  | 10 WAT | 12 WAT | 14 WAT | 16 WAT | 18 WAT |
| GD | 40.00<sup>h</sup> | 62.47<sup>ab</sup> | 65.23<sup>c</sup> | 72.63<sup>ab</sup> | 78.73<sup>cd</sup> |
| GF | 42.73<sup>fg</sup> | 61.27<sup>ab</sup> | 68.83<sup>b</sup> | 72.23<sup>ab</sup> | 77.37<sup>bcd</sup> |
| GS | 43.77<sup>f</sup> | 64.03<sup>a</sup> | 70.07<sup>a</sup> | 76.83<sup>a</sup> | 76.87<sup>bcd</sup> |
| LL | 50.53<sup>a</sup> | 59.77<sup>c</sup> | 64.57<sup>cd</sup> | 70.80<sup>ab</sup> | 79.17<sup>cd</sup> |
| NPK | 45.71<sup>de</sup> | 60.35<sup>b</sup> | 66.30<sup>c</sup> | 73.44<sup>ab</sup> | 64.60<sup>g</sup> |
| PM | 38.24<sup>i</sup> | 46.40<sup>f</sup> | 60.71<sup>e</sup> | 71.62<sup>c</sup> | 76.36<sup>bcd</sup> |
| GD+GF | 45.37<sup>de</sup> | 62.77<sup>ab</sup> | 69.67<sup>b</sup> | 74.40<sup>a</sup> | 81.17<sup>abc</sup> |
| GD + LL | 49.37<sup>ab</sup> | 65.67<sup>a</sup> | 69.97<sup>b</sup> | 75.27<sup>a</sup> | 85.70<sup>ab</sup> |
| GD+ PM | 46.23<sup>d</sup> | 66.37<sup>a</sup> | 69.57<sup>b</sup> | 73.13<sup>ab</sup> | 87.80<sup>a</sup> |
| GD+NPK | 44.73<sup>de</sup> | 62.27<sup>ab</sup> | 68.83<sup>b</sup> | 77.33<sup>a</sup> | 84.50<sup>ab</sup> |
| GD+GS | 42.13<sup>fg</sup> | 61.70<sup>ab</sup> | 71.87<sup>a</sup> | 73.80<sup>ab</sup> | 78.53<sup>cd</sup> |
| LL+GF | 48.70<sup>c</sup> | 53.63<sup>e</sup> | 60.33<sup>e</sup> | 76.17<sup>a</sup> | 75.37<sup>bcd</sup> |
| LL+PM | 47.67<sup>cd</sup> | 53.27<sup>e</sup> | 66.60<sup>c</sup> | 72.37<sup>ab</sup> | 76.73<sup>bcd</sup> |
| LL+NPK | 49.50<sup>ab</sup> | 52.67<sup>e</sup> | 60.03<sup>e</sup> | 67.60<sup>b</sup> | 70.77<sup>f</sup> |
| LL+GS | 48.30<sup>c</sup> | 56.80<sup>d</sup> | 68.33<sup>b</sup> | 74.97<sup>a</sup> | 74.23<sup>e</sup> |
| PM+NPK | 40.57<sup>h</sup> | 54.50<sup>d</sup> | 63.23<sup>cd</sup> | 73.30<sup>ab</sup> | 75.43<sup>bcd</sup> |
| PM+GS | 49.87<sup>ab</sup> | 59.93<sup>c</sup> | 66.37<sup>c</sup> | 71.00<sup>c</sup> | 81.03<sup>abc</sup> |
| CTRL | 16.67<sup>j</sup> | 39.63<sup>g</sup> | 41.23<sup>f</sup> | 44.90<sup>d</sup> | 40.97<sup>h</sup> |

Values are means of five replicates. Means with the same letter in a column are not significantly different (DMRT at $p < 0.05$ ). AMF =Arbuscular mycorrhizal fungi; PM = Poultry manure; NPK= Nitrogen-Phosphorus-Potassium fertilizers; GS = Green Manure of *Gliricidia sepium*; LL: *Leucaena leucocephala*; GD: *Glomus deserticola*; GF; *Glomus fasciculatum* and CTRL= untreated plants.

**Table 2:** Percentage RWC of white Yam treated with AMF, green manure of *G. sepium, Leucaena leucocephala* and other soil amendment on an overcropped soil

| Treatment | Relative water contents (%) |  |  |  |  |
| --- | --- | --- | --- | --- | --- |
|  | 8 WAT | 10 WAT | 12 WAT | 14 WAT | 16 WAT |
| GM | 46.60<sup>c</sup> | 57.70<sup>f</sup> | 64.10<sup>f</sup> | 74.00<sup>b</sup> | 78.10<sup>ab</sup> |
| GE | 49.77<sup>ab</sup> | 59.87<sup>d</sup> | 73.13<sup>b</sup> | 75.13<sup>b</sup> | 78.93<sup>ab</sup> |
| GS | 52.07<sup>a</sup> | 62.27<sup>c</sup> | 72.23<sup>bc</sup> | 79.43<sup>a</sup> | 79.83<sup>ab</sup> |
| LL | 53.00<sup>a</sup> | 58.97<sup>e</sup> | 61.60<sup>g</sup> | 75.20<sup>b</sup> | 76.67<sup>c</sup> |
| NPK | 47.37<sup>c</sup> | 54.73<sup>g</sup> | 50.20<sup>h</sup> | 62.17<sup>hi</sup> | 74.53<sup>d</sup> |
| PM | 52.70<sup>a</sup> | 59.43<sup>d</sup> | 69.93<sup>c</sup> | 75.20<sup>b</sup> | 78.07<sup>ab</sup> |
| GM+GE | 50.73<sup>ab</sup> | 60.43<sup>d</sup> | 70.93<sup>c</sup> | 73.70<sup>c</sup> | 75.13<sup>d</sup> |
| GM + LL | 49.80<sup>ab</sup> | 52.77<sup>h</sup> | 66.33<sup>d</sup> | 71.43<sup>ab</sup> | 79.07<sup>ab</sup> |
| GM+ PM | 45.43<sup>d</sup> | 57.67<sup>f</sup> | 64.57<sup>f</sup> | 68.43<sup>d</sup> | 70.27<sup>f</sup> |
| GM+NPK | 50.73<sup>ab</sup> | 57.90<sup>f</sup> | 66.23<sup>d</sup> | 62.03<sup>ef</sup> | 58.77<sup>g</sup> |
| GM+GS | 52.67<sup>a</sup> | 64.63<sup>b</sup> | 71.93<sup>bc</sup> | 73.00<sup>c</sup> | 73.10<sup>e</sup> |
| LL+GE | 47.93<sup>c</sup> | 59.43<sup>d</sup> | 75.93<sup>a</sup> | 75.00<sup>b</sup> | 82.80<sup>a</sup> |
| LL+PM | 53.70<sup>a</sup> | 66.50<sup>a</sup> | 75.67<sup>a</sup> | 76.20<sup>b</sup> | 76.80<sup>c</sup> |
| LL+NPK | 43.83<sup>e</sup> | 46.67<sup>i</sup> | 50.80<sup>h</sup> | 67.07<sup>d</sup> | 73.30<sup>e</sup> |
| LL+GS | 51.33<sup>ab</sup> | 63.93<sup>bc</sup> | 65.33<sup>e</sup> | 61.33<sup>f</sup> | 77.50<sup>c</sup> |
| PM+NPK | 45.83<sup>d</sup> | 57.13<sup>f</sup> | 67.80<sup>d</sup> | 63.63<sup>e</sup> | 70.47<sup>f</sup> |
| PM+GS | 50.50<sup>ab</sup> | 61.17<sup>c</sup> | 65.67<sup>e</sup> | 67.30<sup>d</sup> | 80.50<sup>b</sup> |
| CTRL | 30.3<sup>f</sup> | 32.50<sup>j</sup> | 35.23<sup>i</sup> | 35.67<sup>g</sup> | 38.17<sup>h</sup> |

Values are means of five replicates. Means with the same letter in a column are not significantly different (DMRT at $p < 0.05$ ). AMF =Arbuscular mycorrhizal fungi; PM = Poultry manure; NPK= Nitrogen-Phosphorus-Potassium fertilizers; GS = Green Manure of *Gliricidia sepium*; LL: *Leucaena leucocephala*; GD: *Glomus deserticola*; GF; *Glomus fasciculatum* and CTRL= untreated plants.

**Table 3:** Correlation coefficient between the growth and yield characters of white yam treated with green manure of *Gliricidia sepium, Leucaena leucocephala*, AMF and other soil amendments

| Parameters | Shoot weight | Root weight | Tuber weight | Tuber Width |
| --- | --- | --- | --- | --- |
| Shoot weight |  |  |  |  |
| Root weight | 0.655** |  |  |  |
| Tuber weight | 0.677** | 0.850** |  |  |
| Tuber width | 0.563** | 0.870** | 0.779** |  |
| Tuber length | 0.732** | 0.843** | 0.748** | 0.839** |

\*\* . Correlation is significant at the 0.01 level (2-tailed).

Furthermore, there was a positive relationship between tuber weight and relative water content (r=0.812) and chlorophyll content (r=0.845) at p> 0.05 level of significant (Table 4).

**Table 4:**
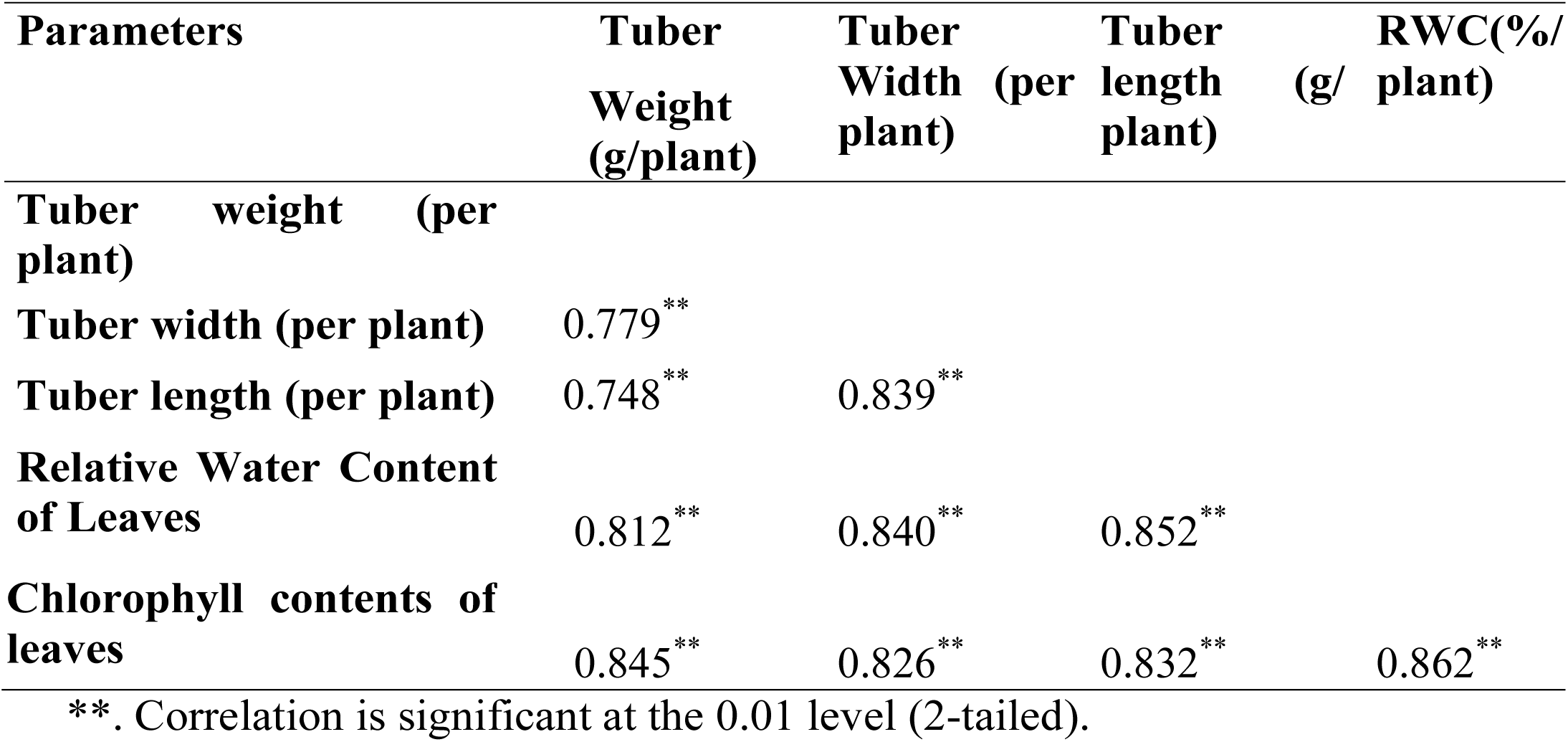
The relationship between the yield and physiological characters of white yam treated with green manure of *Gliricidia sepium, Leucaena leucocephala*, AMF and other soil amendments

| Parameters | Tuber Weight (g/plant) | Tuber Width (per plant) | Tuber length (per plant) | RWC(%/ plant) |
| --- | --- | --- | --- | --- |
| Tuber weight (per plant) |  |  |  |  |
| Tuber width (per plant) | 0.779** |  |  |  |
| Tuber length (per plant) | 0.748** | 0.839** |  |  |
| Relative Water Content of Leaves | 0.812** | 0.840** | 0.852** |  |
| Chlorophyll contents of leaves | 0.845** | 0.826** | 0.832** | 0.862** |

\*\* . Correlation is significant at the 0.01 level (2-tailed).

Table 5 shows the effect of Arbuscular mycorrhizal fungi inoculation, green manure of *Gliricidia sepium* (GS), *Leucaena leucocephala* (LL) and other soil amendments on the growth and yield of white yam. The combined treatment of GS+LL enhanced the higher shoot weight (774.8 g) in white yam, while the untreated had the least (110.2 g). The result shows no significant difference between white yam treated with GS+LL and PM+GS; PM+NPK; GD+NPK; GM+GS and LL treated plants in their shoot weight. White yam treated with poultry manure (PM) had the highest root weight (19.2) which was significantly different from all the other treatments. Many workers opined that green manure (organic fertilizers) increase leaf number and area, shoot weight and productivity of plants (Kaur *et al*., 2005; Benjawan *et al*., 2007; Murmu *et al*., 2013).

**Table 5:** Effects of green manure, Arbuscular mycorrhizal fungi inoculations and other soil amendments on the growth and yield of white yam

| Treatment | Shoot weight (g) | Root weight (g) | Tuber weight (kg) |
| --- | --- | --- | --- |
| GD | 422.31<sup>d</sup> | 19.20<sup>b</sup> | 3.5<sup>e</sup> |
| GF | 391.44<sup>e</sup> | 16.70<sup>e</sup> | 2.9<sup>f</sup> |
| GS | 757.00<sup>a</sup> | 19.50<sup>b</sup> | 2.4<sup>g</sup> |
| LL | 749.61<sup>a</sup> | 16.31<sup>e</sup> | 3.1<sup>f</sup> |
| NPK | 658.03<sup>b</sup> | 17.02<sup>d</sup> | 3.6<sup>e</sup> |
| PM | 733.60<sup>a</sup> | 20.20<sup>a</sup> | 3.8<sup>d</sup> |
| GD+GF | 421.71<sup>d</sup> | 17.21<sup>d</sup> | 4.5<sup>b</sup> |
| GD + LL | 372.26<sup>e</sup> | 18.14<sup>c</sup> | 3.5<sup>d</sup> |
| GD+ PM | 681.60<sup>b</sup> | 19.00<sup>b</sup> | 4.3<sup>c</sup> |
| GD+NPK | 719.33<sup>a</sup> | 17.20<sup>d</sup> | 4.6<sup>b</sup> |
| GD+GS | 746.21<sup>a</sup> | 17.02<sup>d</sup> | 3.9<sup>d</sup> |
| LL+GF | 551030<sup>c</sup> | 18.22<sup>c</sup> | 3.0<sup>f</sup> |
| LL+PM | 536.00<sup>c</sup> | 16.70<sup>e</sup> | 4.1<sup>c</sup> |
| LL+NPK | 680.05<sup>b</sup> | 16.83<sup>e</sup> | 3.7<sup>d</sup> |
| LL+GS | 764.83<sup>a</sup> | 18.32<sup>c</sup> | 4.7<sup>b</sup> |
| PM+NPK | 739.63<sup>a</sup> | 18.00<sup>c</sup> | 4.7<sup>b</sup> |
| PM+GS | 760.41<sup>a</sup> | 18.91<sup>c</sup> | 5.5<sup>a</sup> |
| CTRL | 180.00<sup>f</sup> | 11.00<sup>f</sup> | 1.6<sup>h</sup> |

Values are means of five replicates. Means with the same letter in a column are not significantly different (DMRT at $p < 0.05$ ).
AMF: Arbuscular mycorrhizal fungi; PM: Poultry manure; NPK: Nitrogen-Phosphorus-potassium fertilizers; GS: Green Manure of *Gliricidia sepium*; LL: *Leucaena leucocephala*; GF: *Glomus fasciculatum*; GD *Glomus deserticola* and CTRL= untreated plants.

It was observed that there was no significant difference in the root weight between GD; GS; and GD+PM treated plants. This is because green manure increases photosynthetic activities in plants, this will lead to higher photosynthates assimilated as tuber especially in yams (Bationo *et al*., 1992; Yadana *et al*., 2009). Tuber weight of white yam was enhanced by the combined treatment of plants and animal manure (PM+GS; LL+GS; LL+PM and GD+PM) (Buerkert *et al*., 2001). White yam inoculated with *Glomus deserticola* (GD) and the combined treatment of *Glomus deserticola* and *Glomus fasciculatum* (GD+GF) also enhanced tuber production. There was no significant difference between the tuber yield of GD and NPK treated plants. This result is similar to existing report, that growth and yield of plants were significantly enhanced by AMF and that the shoot dry weight of *G. deserticola* inoculated plants was almost twice of non-mycorrhizal plants (Changxian *et al*., (2008).

## CONCLUSION

African soil is faced with desertification, erosion, deforestation, excessive use of inorganic fertilizers which may be washed into nearby rivers, kill humus and soil microorganisms etc. Sustainable white yam production in this region is dependent on the good maintenance of the ecosystem. Bio-fertilizers and organic manure used in this trial enhanced white yam tubers production, this can conveniently replace NPK and other inorganic fertilizers.

## ACKNOWLEDGMENTS

The author wishes to express his profound appreciation to the Tertiary Education Trust Fund (TETFUND) for the financial assistance.

